# Hexapeptide induces M2 macrophage polarization via the JAK1/STAT6 pathway to promote angiogenesis in bone repair

**DOI:** 10.1101/2021.06.25.449833

**Authors:** Xinyun Han, Junxian Hu, Wenbo Zhao, Hongwei Lu, Jingjin Dai, Qingyi He

## Abstract

Angiogenesis is essential for successful bone defect repair. In normal tissue repair, the physiological inflammatory response is the main regulator of angiogenesis through the activity of macrophages and the cytokines secreted by them. In particular, M2 macrophages which secrete high levels of PDGF-BB are typically considered to promote angiogenesis. A hexapeptide [WKYMVm, (Trp-Lys-Tyr-Met-Val-D-Met-NH2)] has been reported to modulate inflammatory activities. However, the underlying mechanisms by which WKYMVm regulates macrophages remain unclear. In this study, the possible involvement by which WKYMVm induces the polarization of macrophages and affects their behaviors was evaluated. In vitro results showed that macrophages were induced to an M2 rather than M1 phenotype and the M2 phenotype was enhanced by WKYMVm through activation of the JAK1/STAT6 signaling pathway. It was also found that WKYMVm played an important role in the PDGF-BB production increase and proangiogenic abilities in M2 macrophages. Consistent with the results in vitro, the elevated M2/M0 ratio induced by WKYMVm enhanced the formation of new blood vessels in a femoral defect mouse model. In summary, these findings suggest that WKYMVm could be a promising alternative strategy for angiogenesis in bone repair by inducing M2 macrophage polarization.

## 1. Introduction

Bone defects caused by trauma, infection, cancer, congenital diseases and other factors are common diseases endangering human health (1–3). Bone repair remains a great challenge in orthopedics because of poor vascularization. Vascularization is essential for bone defect repair, and the degree of bone repair is directly proportional to the degree of vascularization at the bone defect site (4, 5). Therefore, vascularization is a basic research problem that needs to be solved urgently.

Macrophages are the key regulators of inflammation-related immunity and have multiple functions (e.g., angiogenesis and injury repair) (6, 7). Importantly, macrophages have excellent plasticity, and they may respond to various stimuli effectively via their polarization into different phenotypes (8, 9). Classic macrophages (M1) are activated by microbial agents (lipopolysaccharides and interferon-γ) that have proinflammatory effects. M1 macrophages secrete major inflammatory cytokines, including interleukin (IL)-1β and tumor necrosis factor (TNF)-α, and upregulate enzyme-inducible nitric oxide synthase (iNOS) (10, 11). The other type of macrophages (M2) are alternately activated by IL-4 or IL-13 and having anti-inflammatory effects. M2 macrophages upregulate the enzymes arginase (Arg)-1 and chitinase-like protein 3 (Chil3/Ym1) and produce cytokines, including IL-10 and transforming growth factor (TGF)-β (12–14). It is worth noting that M2 macrophages can also promote angiogenesis by secreting high levels of platelet-derived growth factor-BB (PDGF-BB) (15).

The proangiogenic properties of macrophages are not shared among all subgroups because M2 macrophages have been shown to secrete more angiogenic growth factors and cytokines than M1 macrophages do in vitro (16). Although it has been reported that M1 macrophages can also produce proangiogenic cytokines, such as vascular endothelial growth factor (VEGF), to promote angiogenesis, the blood vessels formed only by stimulation with VEGF are immature, leak, and are easily degraded (17). PDGF-BB expressed and secreted by M2 macrophages can recruit pericytes and mesenchymal stem cells to remodel the initial vascular buds, stabilize neovessels, and coordinate osteoblast differentiation (18, 19). The role of M2 macrophages in promoting angiogenesis was first evident in the occurrence and development of malignant tumors. It has been widely recognized that tumor-associated macrophages (TAMs) promote tumor angiogenesis and the formation of vessels at the tumor site (20, 21). The above studies show that M2 macrophages are clearly the prevalent phenotype for angiogenesis during bone repair. Therefore, the enhancement of the M2 phenotype is considered a means to promote vascularization. However, the role of nontumor-associated M2 macrophages in angiogenesis and their production are currently poorly understood.

M2 macrophages are induced by activating various signaling pathways. Signal transducer and activator of transcription (STAT) 3, STAT6, interferon regulatory factor (IRF) 4, and peroxisome proliferator-activated receptor ◻ (PPAR◻) are important transcription factors involved in M2 polarization (22–24). In addition, IL-4/13 is a cytokine known to induce polarization of M2 macrophages through the Janus kinase (JAK)1/STAT6 pathway, which is of great significance to tissue repair, elimination of apoptotic cells, and inhibition of inflammatory responses (25, 26). IL-4 activates JAK phosphorylation by binding to IL-4Rα on the membrane, and JAK further phosphorylates STAT6, allowing it to enter the nucleus. After nuclear entry, STAT6 binds to the nuclear receptor PPARγ to mediate M2 polarization (27). This pathway is a key signaling node for the activation of M2 macrophages (26).

WKYMVm (Trp-Lys-Tyr-Met-Val-D-Met-NH2) is a modified hexapeptide identified by screening synthetic peptide libraries (28). Formyl peptide receptor (FPR), which belongs to the G protein-coupled receptor family, is a known high-affinity receptor for WKYMVm. FPR is mainly expressed in phagocytic cells, such as neutrophils, monocytes, and macrophages, and plays an important role in host defense and inflammation. It has three subtypes, namely, FPR1, FPR2/ALX, and FPR3 (28). It is interesting that WKYMVm, by binding to FPR2, shows immunomodulatory capabilities, especially immunosuppressive activity, under different physiological and pathological conditions (29, 30). This process is initiated by FPR2 interacting with agonists to activate other signaling molecules, such as JAK, phosphoinositide 3-kinase (PI3K), protein kinase B (AKT), and mitogen-activated protein kinases (P38, ERK, and JNK), thus ultimately promoting cell migration, phagocytosis, and inflammatory cytokine secretion (31, 32). However, the effect of WKYMVm on macrophages, key immune regulators, has hardly been studied, and the underlying mechanism is still unclear. Studies have found that in human lung cancer cells, CaLu-6, WKYMVm can bind to FPR2 to mediate epidermal growth factor receptor (EGFR) transactivation and further trigger the JAK1/STAT6 pathway (28). Therefore, it is reasonable to speculate that WKYMVm may modulate macrophages through the JAK1/STAT6 pathway.

In this study, it was hypothesized that WKYMVm can polarize macrophages to the M2 type, thus indirectly affecting remodeling and forming new blood vessels for bone repair. The results from this work show that WKYMVm induces polarization of M2 macrophages in vivo and in vitro, leading to angiogenesis. Additionally, the molecular mechanism underlying these phenomena was found to be based on the activation of the JAK1/STAT6 pathway mediated by WKYMVm. This study suggests that WKYMVm exerts a regulatory effect on the macrophage phenotype for angiogenesis, thus providing new ideas for promoting vascularization in the repair of bone defects.

## 2. Materials and Methods

### 2.1. Isolation and culture of mouse bone marrow macrophages (BMMs)

After euthanasia, bone marrow cells were isolated from the tibias and femurs of 4-week-old C57 mice, and they were resuspended in red blood cell (RBC) lysate. The collected cells were cultured in complete alpha minimum essential medium (α-MEM; Gibco, USA) containing 1% penicillin-streptomycin (Thermo Fisher Scientific), 10% fetal bovine serum (FBS; Gibco, USA), and 30 ng/ml macrophage colony-stimulating factor (M-CSF; R&D Systems, USA) at 37°C and 5% CO2 for 3 days. The attached cells were considered BMMs in which proliferation was consistently stimulated by 30 ng/ml M-CSF.

### 2.2. Cell culture and preparation of conditioned media

RAW264.7 cells (mouse macrophage cells) were obtained from the American Type Culture Collection (ATCC, USA) and cultured in Dulbecco’s modified Eagle’s medium (DMEM; Gibco, USA) supplemented with 1% penicillin-streptomycin and 10% FBS at 37°C and 5% CO2. For collection of macrophage-conditioned medium, BMMs were pretreated for 24 h with complete α-MEM supplemented with different concentrations of WKYMVm, which was synthesized by GL Biochem (Chengdu, China) with >95% purity. The cells were then thoroughly washed and cultured in new normal complete α-MEM medium for 24 h at 37°C and 5% CO2 to eliminate direct interference from WKYMVm. The supernatant from BMMs was obtained as conditioned medium and used in subsequent lumen formation and scratch tests.

### 2.3. Cytotoxicity assay

RAW264.7 cells (3×103 cells/well) or BMMs (5×103 cells/well) were inoculated into two 96-well plates in triplicate and then treated with different concentrations of WKYMVm (0, 0.1, 1, 5, 10, and 20 μM) for 24 h and 48 h. Cell viability was tested using the Cell Counting Kit-8 (CCK-8; Beyotime, Beijing, China) assay according to the manufacturer’s instructions. After the specified time, fresh medium containing 1/10 CCK-8 reagent was then added to each well and incubated at 37°C for 1 h. The absorbance of all wells was measured at 450 nm using a 96-well plate reader (Bio-tek, Synergy, USA).

### 2.4. RNA isolation and real-time quantitative PCR (qPCR)

Total RNA was isolated using TRIzol reagent (Thermo Fisher Scientific), and total RNA was reverse-transcribed into cDNA by a reverse transcription kit (Takara Bio Inc., Japan). Then, real-time qPCR was performed using SYBR premix ex-Taq-ii (Takara Bio Inc., Japan) and a PCR detection system (Bio-Rad, Hercules, CA, USA). GAPDH levels were used as the internal standardization control for mRNA levels. The thermal cycles were as follows: first predenaturation step at 95°C for 30 s, then the samples were amplified 40 times at 95°C for 5 s, at 60°C for 1 min, and finally at 65°C for 5 s. The list of primers is shown in Table 1.

**Table 1.**
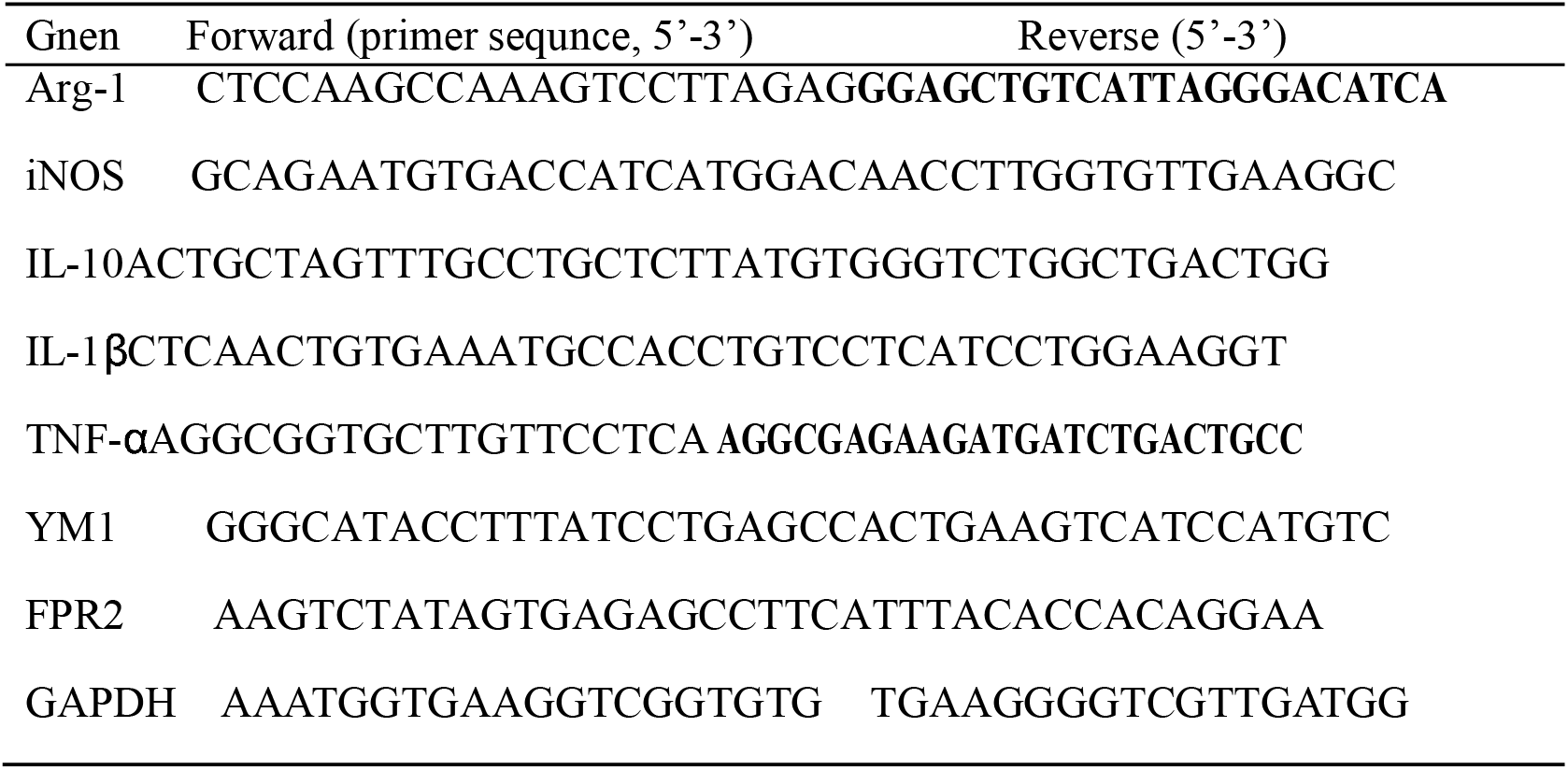
Primer sequences for qPCR

### 2.5. Western blot

After the assigned treatment, cells were lysed in cell lysis buffer containing PMSF supplemented with a protease inhibitor or protease and phosphatase inhibitor cocktail. The proteins were separated using gel electrophoresis and electrotransferred to polyvinylidene fluoride membranes. After blocking with 5% bovine serum albumin for 2 h and washing with Tris-buffered saline/Tween-20 (TBS-T) for 30 min, the membranes were incubated overnight at 4°C with primary antibodies against Arg-1 (1:1000), CD206 (1:1000), CD86 (1:1000), iNOS (1:1000), PPARγ (1:1000) (Bioworld, USA), JAK1 (1:1000), STAT6 (1:1000), p-JAK1 (1:1000), p-STAT6 (1:1000) (Cell Signaling Technology, USA), β-tubulin (1:5000) or β-actin (1:5000) (Bioworld, USA). Subsequently, the membranes were washed with TBS-T for 30 min at room temperature and incubated with horseradish peroxidase-conjugated secondary antibodies for 2 h. The membranes were then washed three times with TBS-T. Images of protein bands were obtained using the Chemi Doc XRS+ Imaging System (Bio-Rad), and the densitometry values were analyzed by ImageJ. β-Actin or β-tubulin served as an internal control.

### 2.6. Flow cytometry

BMMs (1×106) cultured with WKYMVm for 24 h were collected in 1.5 ml Ep tubes after washing twice and were incubated with Alexa Fluor-488 (AF488, BioLegend, USA)-conjugated anti-mouse CD206 and activated protein C (APC, BioLegend, USA)-conjugated anti-mouse CD86 at room temperature for 1 h without light. Finally, cells were suspended in 150 μl PBS and then analyzed by FACSCalibur flow cytometry (BD Biosciences, USA).

### 2.7. Tube formation assay

HUVECs (human umbilical vein endothelial cells) from ATCC (Rockville, MD, USA) were seeded at a concentration of 5× 104 cells/well onto Matrigel (Biosciences, USA)-coated 24-well plates and then treated with the above conditioned medium from BMMs prestimulated with WKYMVm or not. After 8 h, tube formation was imaged by an inverted microscope (Leica DMI6000B, Germany).

### 2.8. Migration assay

For the scratch wound migration assay, HUVECs seeded at 6-well plates to confluence were scratched using a 1 ml pipette. Then, they were treated with conditioned media of BMMs pretreated with WKYMVm or not. At 0 h, 12 h, and 24 h post wounding, cells were imaged, and migration was measured by the relative area of cell coverage from the wound edges.

### 2.9. Immunofluorescence

For cell assays, BMMs plated on 96-well plates were washed, fixed, and permeabilized with 4% paraformaldehyde (0.3% Triton X-100). After 30 min of blocking with 1% BSA, the cells were incubated overnight at 4◻°C with primary antibodies against CD206 and CD86 (1:100; Abcam, UK). On the second day, cells were incubated with secondary antibodies conjugated with AF488 and Cy3 (Abcam, UK) for 1◻h at room temperature. Finally, the nucleus was stained with DAPI for 5 min.

For bone tissue assays, femurs were dissected and then fixed in 4% paraformaldehyde at 4◻°C for 48 h. Subsequently, femurs were embedded in paraffin after decalcification in 10% EDTA for two weeks. Samples were longitudinally cut into 4-μm-thick sections along the sagittal plane of the femur, including the metaphysis and diaphysis. Then, bone sections were double stained with AF647 anti-mouse F4/80 and AF488 anti-mouse CD206 (BioLegend, USA) overnight at 4°C while avoiding light.

### 2.10. ELISA

The supernatants of BMM cultures treated with WKYMVm for 24 h and 48 h were collected. According to the instructions of the ELISA kit (Novus USA), the levels of PDGF-BB and VEGF secreted by cells were assessed with 2 pg/ml sensitivity. Each sample was set up in three wells. Each measurement was calculated by a dilution calibration curve from the recombinant mouse standard.

### 2.11. Immunohistochemical analyses

Each bone tissue section, as described previously, was stained for CD31 (Bioworld Technology, MN, USA) and assessed by immunohistochemistry (IHC). The images were obtained by full fragment scanning.

### 2.12. Femoral defect model in mice

A 1-mm diameter hole was made bilaterally on femoral condyles of 30 adult C57 mice anesthetized with 10% pentobarbital, and gelatin sponges soaked in WKYMVm (100 μM) or PBS, respectively were implanted into the defect sites. The mice were locally injected with WKYMVm or an equal volume of PBS 3 times a week. To efficiently deplete macrophages in mouse bone marrow, a standard macrophage depletion kit (Encapsula NanoScience, USA) was used by intravenous injection and administered continuously according to the instructions (33, 34). Briefly, mice were randomly divided into three groups: control (PBS), WKYMVm and WKYMVm+clodronate liposomes. Animals were sacrificed one week, two weeks or 6 weeks after the operation, and femurs were harvested for immunohistochemistry, immunofluorescence and micro-CT analysis. The entire animal study was performed in accordance with the guidelines of the animal protection and use committee of Army Medical University.

### 2.13. Statistical analysis

Data are expressed as the mean◻±◻SD (n≥3). Student’s tests were performed to analyze the differences between two groups using SPSS 22.0 software. p < 0.05 indicates statistical significance.

## 3. Results

### 3.1 Toxicity evaluation of WKYMVm in RAW264.7 cells and BMMs

RAW264.7 cells and BMMs were treated with a concentration gradient of WKYMVm (Figure 1A) for 24 h and 48 h, respectively, followed by the CCK8 reagent to estimate cell viability. The results revealed that WKYMVm at concentrations ≤20 μM did not affect cell proliferation (Figure 1B, C). Therefore, WKYMVm concentration gradients of 0.1 μM, 1 μM, and 10 μM were used in further studies.

**Figure 1.**
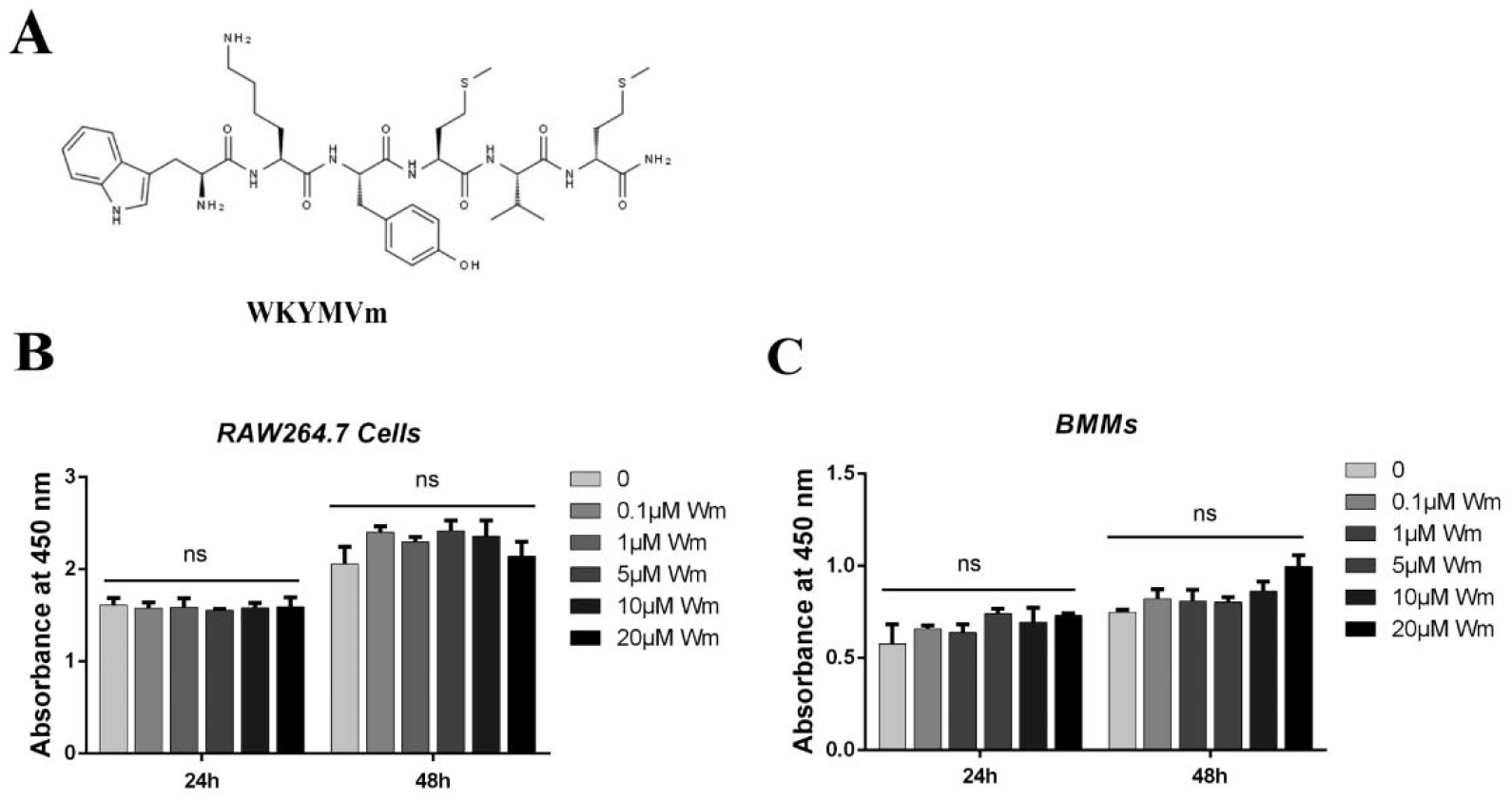
Toxicity evaluation of WKYMVm in RAW264.7 cells and bone marrow macrophages (BMMs). A, Chemical structure of WKYMVm. B and C, Cell viability assay of RAW264.7 cells or BMMs after WKYMVm (0, 0.1, 1, 5, 10, and 20 μM) treatment for 24 h or 48 h, as determined by the CCK8 assay. Data represent the means ± SD (n=3). *p < 0.05; **p < 0.01; and ***p < 0.001; ns, not significant relative to the control (0 μM).

### 3.2 WKYMVm induces M2 macrophage polarization in vitro

To study the effect of WKYMVm on macrophages in vitro, BMMs were isolated. We assumed that M-CSF-induced BMMs and RAW264.7 cells were M0 macrophages. After 24 h of WKYMVm stimulation, CD206+ macrophages (M2) were significantly increased with an increase in the WKYMVm concentration by up to 50.9%, as demonstrated by flow cytometry analysis. However, there was no marked difference in CD86+ macrophages (M1) in the WKYMVm-induced groups (including 0 μM), and the average percentage was only 2.03% (Figure 2A, B). The results of cell immunofluorescence similarly showed that CD206 (M2-specific marker) was more highly expressed after WKYMVm treatment, but CD86 (M1-specific marker) was still barely expressed (Figure 2C). Based on these results, we initially observed that M0 macrophages were induced to M2 macrophages rather than M1 macrophages by WKYMVm.

**Figure 2.**
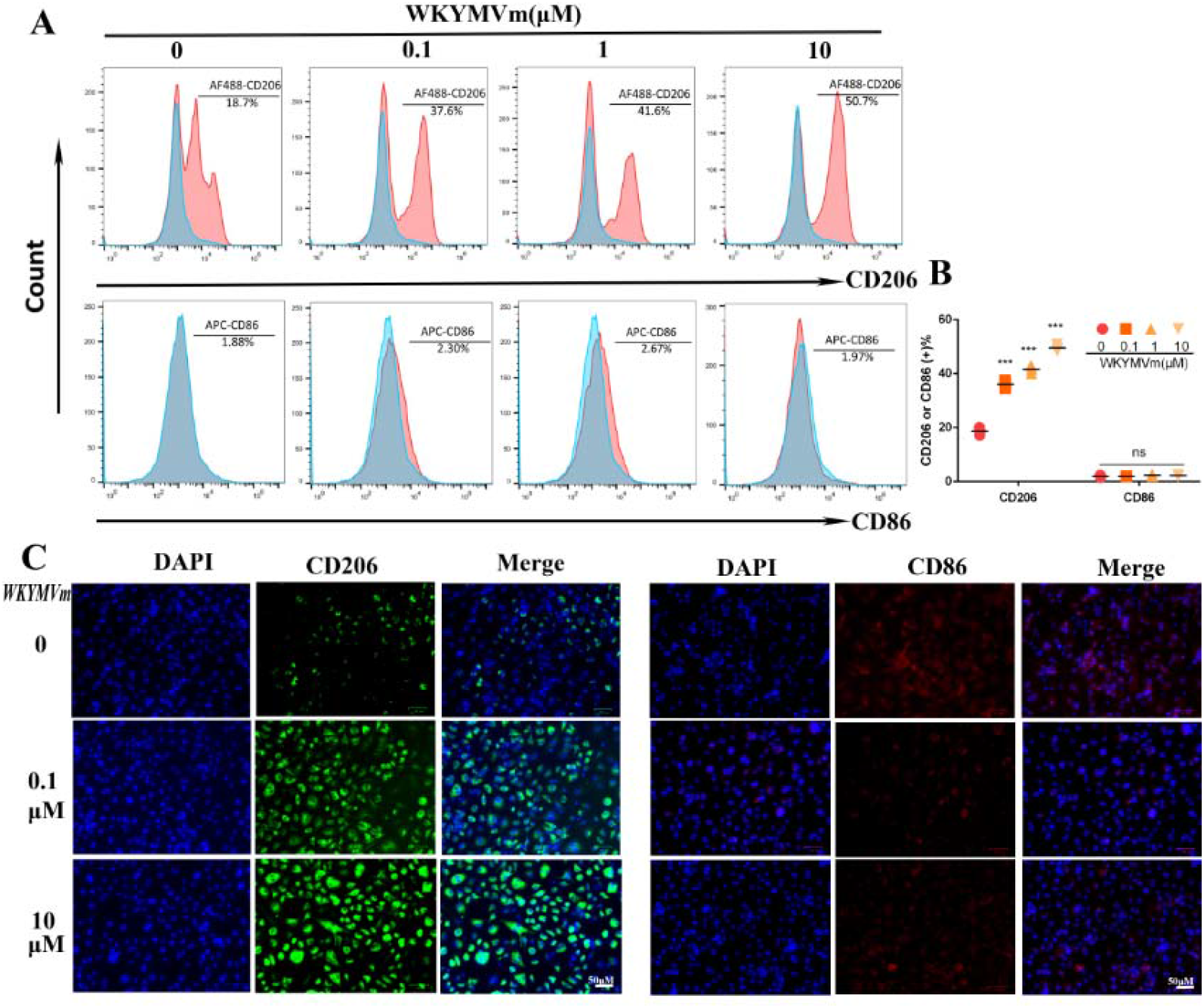
WKYMVm induced polarization of M0 macrophages into M2 macrophages. A, Surface marker CD206 (M1) and CD86 (M2) expression in BMMs treated with WKYMVm (0.1, 1, and 10 μM) for 24 h was measured by flow cytometry. Representative flow cytometry images are shown. B, Quantification of CD206 and CD86 expression in (A). Data represent the means ± SD (n=3). *p < 0.05; **p < 0.01; and ***p < 0.001; ns, not significant relative to the control (0 μM). C, Immunostaining of CD206 (green) and CD86 (red) in BMMs after WKYMVm (0.1 and 10 μM) treatment for 24 h (×200). Scale bar: 50 μm.

To verify this, we further tested the expression of M2 markers (Arg-1, YM1, and IL-10) and M1 markers (iNOS, TNF-α, and IL-1β). The mRNA expression levels of Arg-1, YM1, and IL-10 in WKYMVm-treated BMMs were significantly increased, whereas those of iNOS, TNF-α, and IL-1β were decreased (Figure 3A). Western blot results both in RAW264.7 cells and BMMs revealed that protein expression was consistent with the results of mRNA expression, but iNOS and CD86 were downregulated at WKYMVm concentrations of 1 μM and 10 μM (Figure 3B-E). Taken together, these data indicate that WKYMVm enhanced M2 macrophages and inhibited the M1 macrophage phenotype.

**Figure 3.**
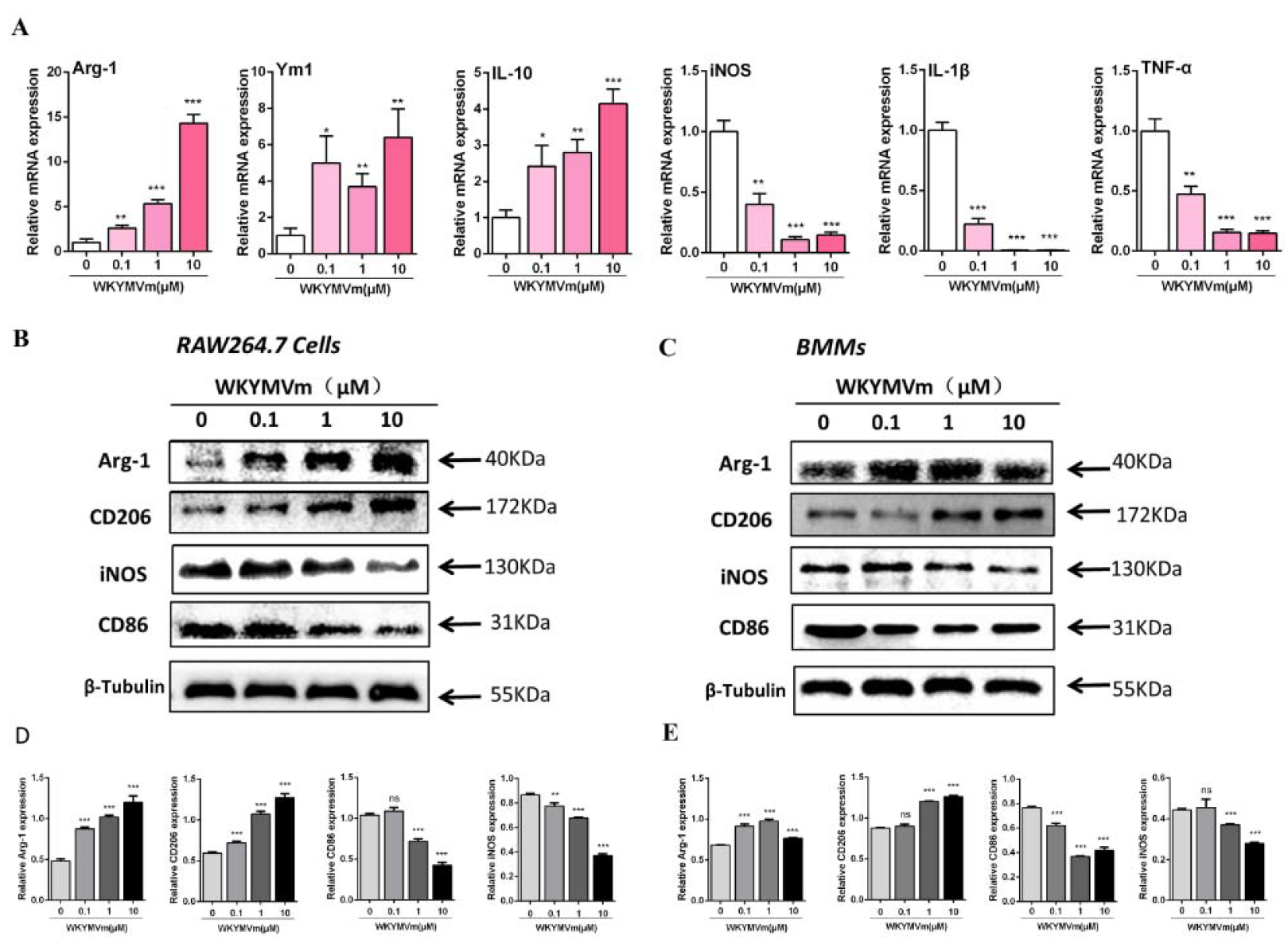
Quantitative polymerase chain reaction (qPCR) and Western blot analysis of M1 and M2 markers for verification. A, BMMs were induced with gradient doses of WKYMVm for 24 h. The expression levels of M2 marker genes (Arg-1, YM1, and IL-10) and M1 marker genes (iNOS, TNF-α, and IL-1β) were analyzed by real-time qPCR (n = 3) B, C, Western blot analysis of marker proteins of M2 and M1 macrophages in RAW 264.7 cells (B) and BMMs (C) treated with WKYMVm for 48 h. β-Tubulin was used as a loading control. D, E, Quantification of the expression of marker proteins in (B) and (C) was performed using ImageJ. Data represent the means ± SD (n=3). *p < 0.05; **p < 0.01; and ***p < 0.001; ns, not significant relative to the control (0 μM).

### 3.3 WKYMVm enhances the proangiogenic effect of M2 macrophages

The above findings suggest that WKYMVm induces the polarization of M2 macrophages. To determine the effect of WKYMVm-induced M2 macrophages on angiogenesis, we examined the expression of the PDGF-BB gene in BMMs and RAW264.7 cells induced by WKYMVm, and the results showed that PDGF-BB expression was upregulated in both cell lines (Figure 4A). Subsequently, we used ELISA to detect the secretion of PDGF-BB and VEGF in BMMs and showed that compared with that in the control group, PDGF-BB and VEGF secretion were enhanced by WKYMVm, and this difference was most significant in the 24 h-induced cells. However, VEGF secretion stopped increasing after 10 μM WKYMVm treatment for 48 h (Figure 4B).

**Figure 4.**
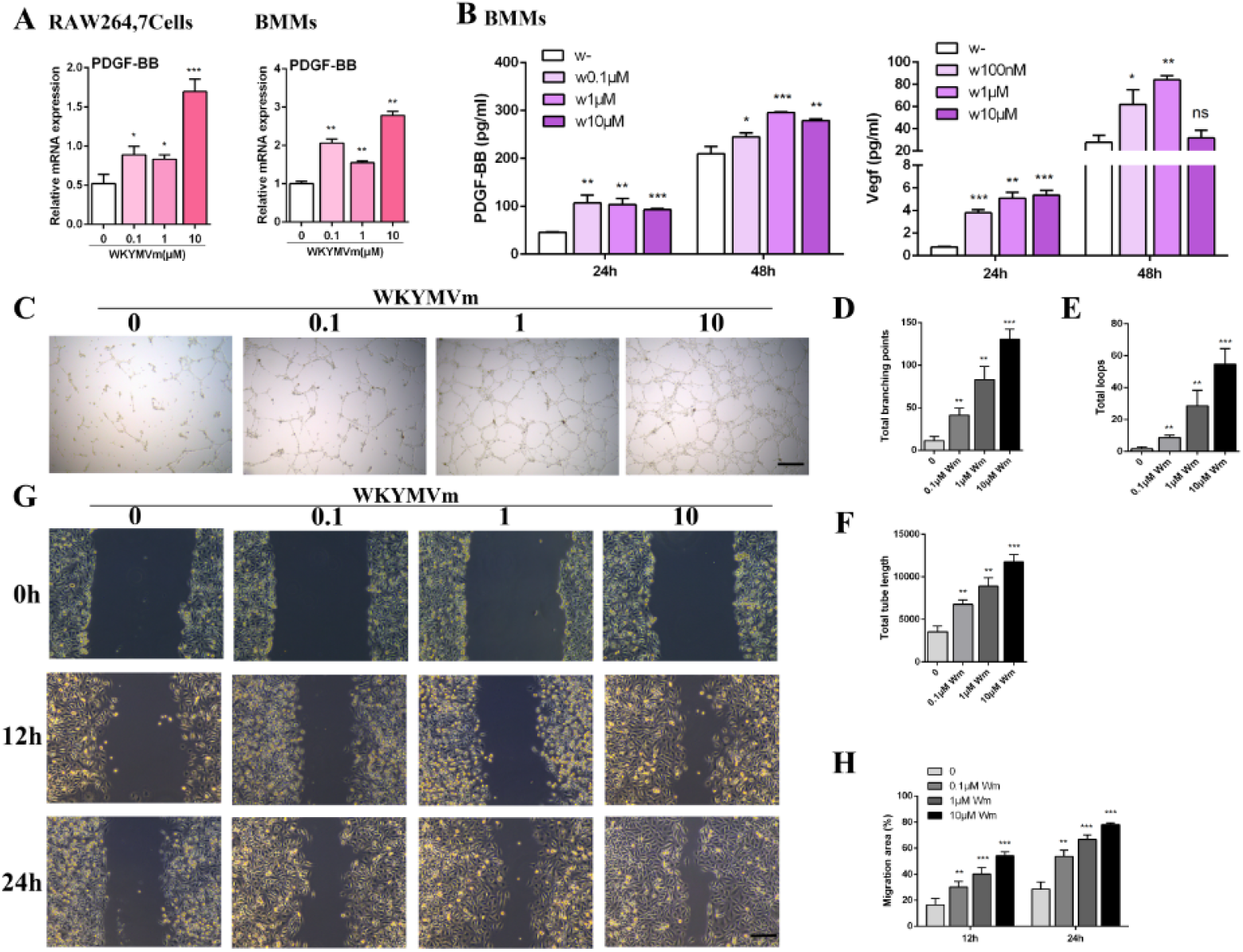
WKYMVm enhances the proangiogenic effect of M2 macrophages in vitro. A, The mRNA expression of PDGF-BB in RAW264.7 cells and BMMs induced by WKYMVm (0.1, 1, and 10 μM) was assessed by real-time qPCR. B, The production of PDGF-BB and VEGF in the cell culture supernatants of WKYMVm-induced macrophages was determined using ELISA. C, Representative images of tube formation assay on Matrigel in HUVECs stimulated with conditioned medium from BMMs in normal medium for 24 h after pretreatment with WKYMVm (0.1, 1, and 10 μM) for 24 h. Scale bar: 100 μm. D-F, Quantitative analyses of the total branching points, total loops and total tube length in (C). G, The migration of HUVECs receiving different treatments (same as C) was detected by the scratch wound assay. Scale bar: 100 μm. H, Quantitative analyses of the migration area in (G). Data represent the means ± SD (n=3). *p < 0.05; **p < 0.01; and ***p < 0.001; ns, not significant relative to the control (0 μM).

Furthermore, we performed a series of angiogenesis-related assays in HUVECs. The lumen formation test on Matrigel is an in vitro angiogenesis model, and the measurements of total tube length, total branch points, and total loops (Figure 4C-E) can be used to quantify the ability of HUVECs to form vessels. As shown in Figure 4B, all HUVEC parameters were increased under the stimulation of WKYMVm-treated BMM-conditioned medium, and they showed the strongest vessel-forming ability at 10 μM WKYMVm. The scratch test revealed that WKYMVm-induced conditioned media could significantly increase HUVEC motility compared to the control; when exposed to conditioned medium of macrophages stimulated with 10 μM WKYMVm, the migration of HUVECs was the greatest (Figure 4F, G). Generally, these tests showed that WKYMVm-induced M2 macrophages have an enhanced angiogenic ability.

### 3.4 WKYMVm induces M2 polarization through the JAK1/STAT6 pathway

We focused on JAK1/STAT6 signaling downstream of the WKYMVm receptor FPR2, which is necessary for M2 polarization. First, the expression of FPR2 was increased by WKYMVm (Figure 5A). To perform the assays, BMMs were stimulated with PBS or WKYMVm (10 μM) for 0, 15, 30, or 60 min after starvation in serum-free DMEM for 12 h. The results showed that the levels of JAK1 and STAT6 protein remained unchanged, while WKYMVm significantly stimulated the phosphorylation of JAK1 and STAT6 at 30 min compared to the control treatment (Figure 5B). Statistical analysis showed that the relative levels of p-JAK1/JAK1 and p-STAT6/STAT6 in WKYMVm-treated cells were significantly higher than those in untreated cells (Figure 5C). PPARγ is a nuclear transcription factor, and its activation can enhance the polarization of circulating monocytes to form M2 macrophages. Therefore, we further determined that the expression of PPARγ was significantly increased in BMMs after stimulation with WKYMVm (10 μM) for 24 h (Figure 5D, E). These results indicate that the JAK1/STAT6 signaling pathway is significantly activated.

**Figure 5.**
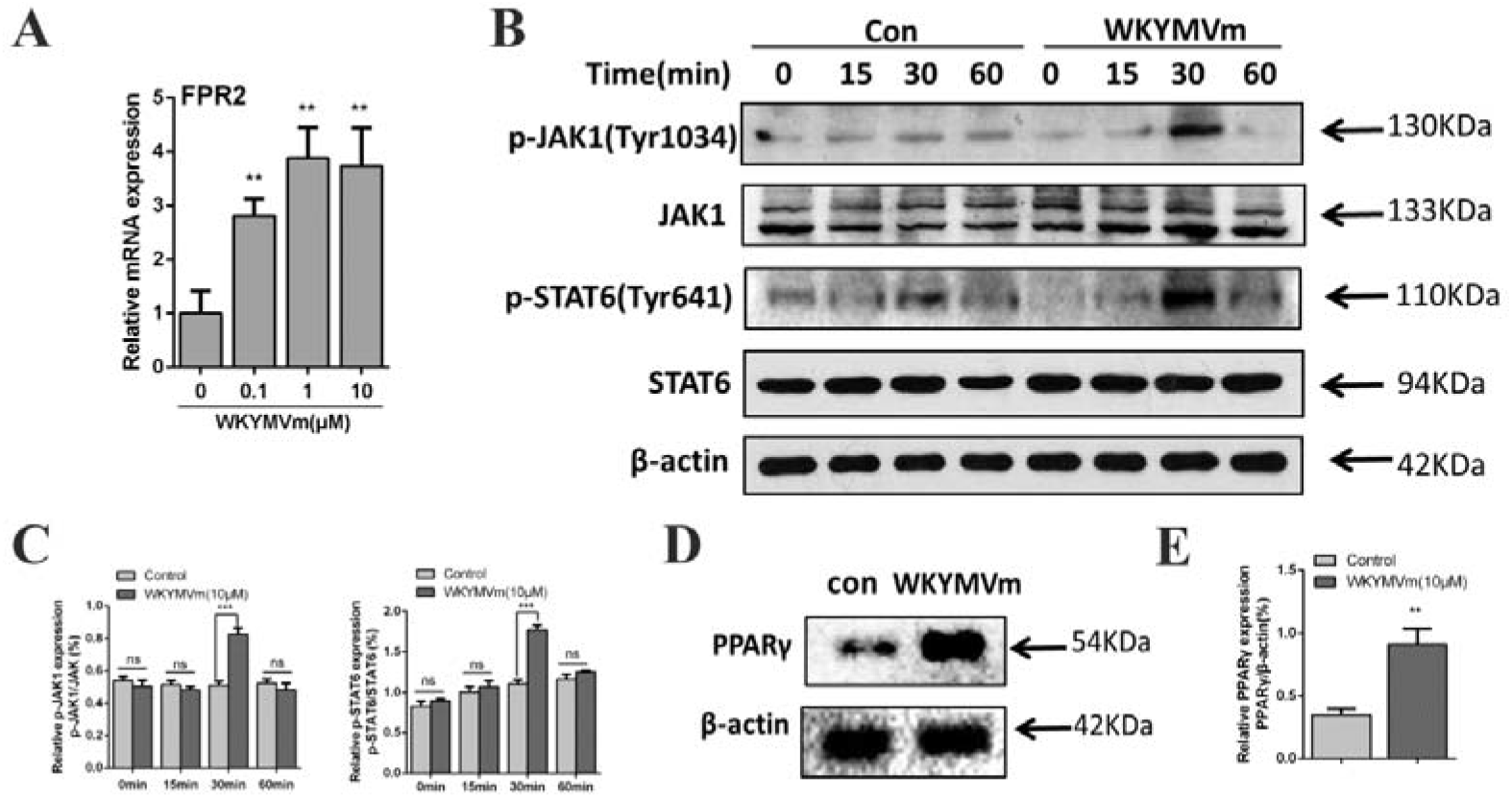
WKYMVm induces M2 polarization through the JAK1/STAT6 pathway. A, The mRNA expression of FPR2 in BMMs was assessed by real-time qPCR. B, Phosphorylation levels of JAK1 and STAT6 in BMMs after WKYMVm (0 or 10 μM) treatment for 0, 15, 30, or 60 min by Western blot. β-Actin was used as a loading control. C Quantification of p-JAK1 and p-STAT6 relative to total JAK1 and STAT6. D. BMMs were incubated with or without WKYMVm for 48 h. The expression of PPAR◻ was analyzed using Western blot. E, Quantification of PPARγ expression relative to β-tubulin expression. Data represent the means ± SD (n=3). *p < 0.05; **p < 0.01; and ***p < 0.001; ns, not significant relative to the control (0 μM).

### 3.5 JAK1 inhibitor ruxolitinib prevents M2 polarization and inhibits angiogenesis

To block the JAK1/STAT6 pathway, we used ruxolitinib, a small molecule that selectively and reversibly inhibits the JAK1 and JAK2 enzymes. BMMs were treated with or without ruxolitinib (2 μM) for 2 h before being treated with PBS or WKYMVm (10 μM) for 24 h. The results from Western blotting (Figure 6A, B) and immunofluorescence (Figure 6C) showed that when cells were treated with ruxolitinib, the increase in Arg-1 and CD206 induced by WKYMVm was reversed. Ruxolitinib inhibited the secretion of PDGF-BB and VEGF from WKYMVm-induced macrophages (Figure 6D), and the ability of HUVECs to induce lumen formation and to migrate in macrophage-conditioned medium from cells induced by WKYMVm was also neutralized (Figure 6E-H). Taken together, these data indicate that WKYMVm induces M2 macrophages to promote angiogenesis through the JAK1/STAT6 pathway. Once the pathway is blocked, M2 cells will not be polarized, and they will lose their angiogenic function.

**Figure 6.**
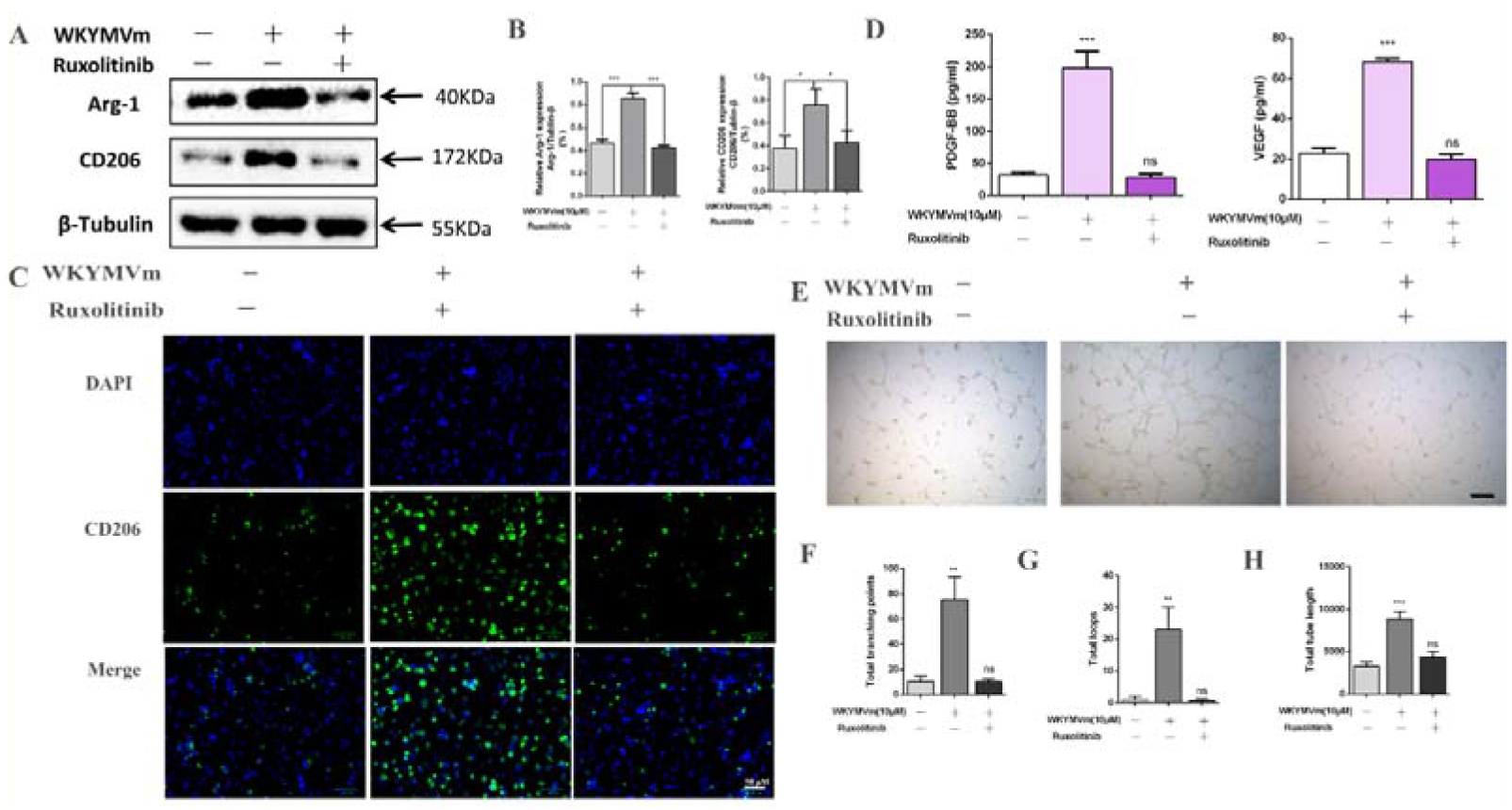
The JAK1 inhibitor ruxolitinib prevents M2 polarization and promotes angiogenesis. A, Western blot analysis of Arg-1 and CD206 (M2) in BMMs treated with PBS, WKYMVm or WKYMVm+ruxolitinib. B, Quantification of expression in (A). C, Expression of CD206 in BMMs with different treatments was analyzed by immunofluorescence assay (200×). D, PDGF-BB and VEGF concentrations in WKYMVm-induced macrophages treated with or without ruxolitinib were determined using ELISA. E, Representative images of tube formation assay on Matrigel in HUVECs stimulated with conditioned media from BMMs in different groups. Scale bar: 100 μm. F-H, Quantitative analyses of total branching points, total loops and total tube length in (E). Data represent the means ± SD (n=3). *p < 0.05; **p < 0.01; and ***p < 0.001; ns, not significant.

### 3.6 WKYMVm-induced M2 macrophages promote angiogenesis, accelerating bone regeneration in vivo

In vivo, a bone defect model in mice was established (Figure 7A).Fourteen days later, as evidenced by IHC staining for CD31 (a vascular endothelial marker), the number of CD31+ blood vessels was significantly increased at the intervention site of femoral condyles in WKYMVm–treated mice compared to PBS-treated mice. In addition, the administration of clodronate liposomes (Standard Macrophage Depletion kit) (33, 34) to WKYMVm–treated mice dramatically reduced blood microvessel generation to a similar extent as that in PBS mice (Figure 7B). This finding showed that WKYMVm-induced macrophages have an important role in angiogenesis in vivo. To identify the polarizations of macrophages with angiogenic functions, F4/80 and CD206 were selected as the pan-macrophage (35) and M2 macrophage markers and were detected by immunofluorescence (IF). The number of F4/80+ cells was significantly reduced in mice upon WKYMVm+ clodronate liposome treatment for 7 and 14 days, confirming that macrophages were successfully depleted (Figure 7C). Interestingly, the expression of F4/80 in WKYMVm–treated mice was similar to that in PBS-treated mice, while CD206 expression was significantly higher (Figure 7C). Combining the immunohistochemical and immunofluorescence results, we concluded that WKYMVm-induced M2 macrophage polarization enhances angiogenesis, consistent with the phenomena in vitro.

**Figure 7.**
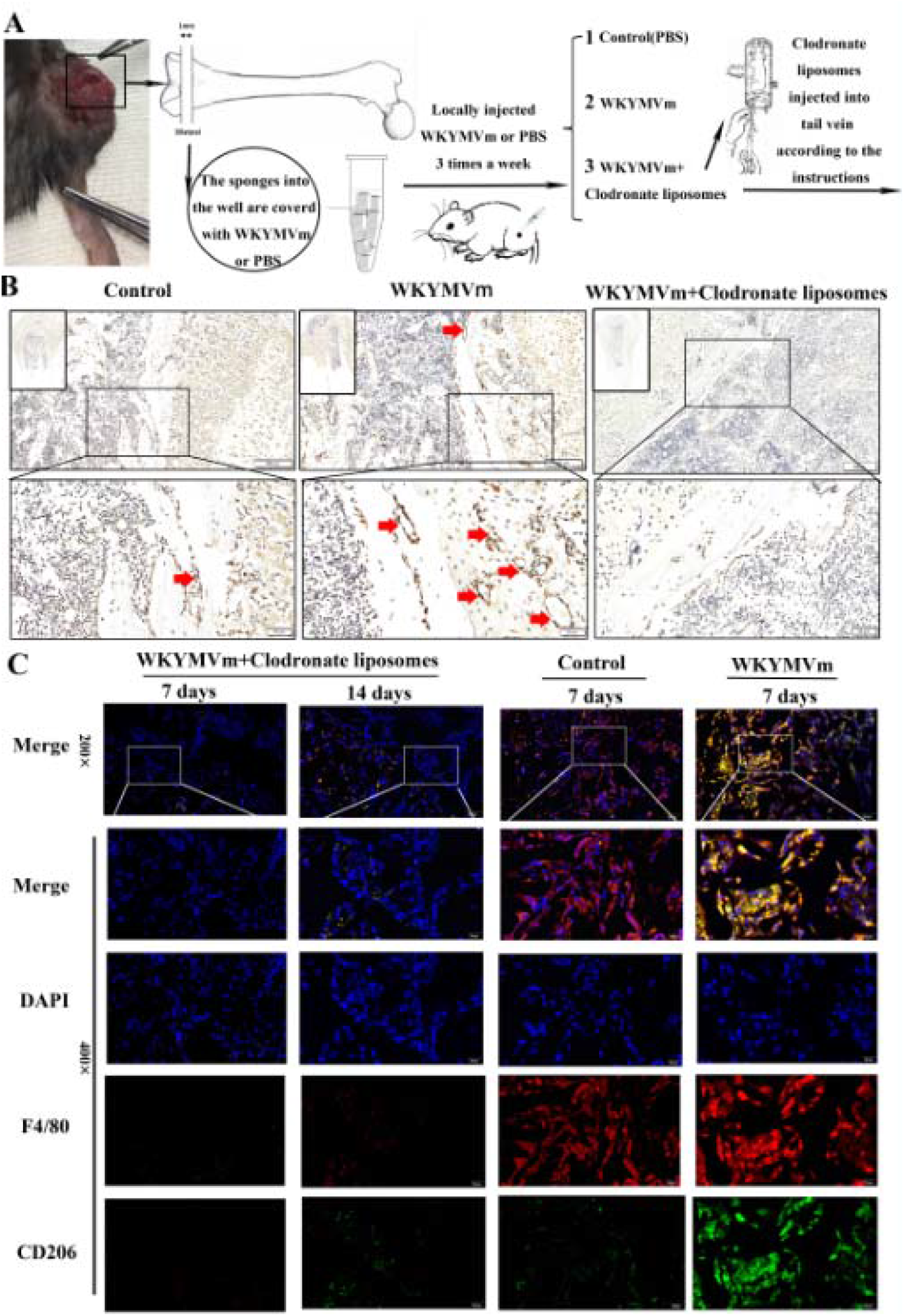
WKYMVm-induced M2 macrophages can promote angiogenesis at bone defect sites in vivo. A, Schematic diagram of the animal study. B, Immunohistochemistry (IHC) of CD31 showed new blood microvessel density (red arrows) at the intervention site of femoral condyles from WKYMVm, WKYMVm+clodronate liposome and control mice after 14 days. C, Immunostaining of F4/80 (red), which represented pan-macrophages, and CD206 (green), which represented M2 macrophages, in distal femur sections from different groups after 7 or 14 days.

Angiogenesis at the bone defect site ultimately aids in bone repair; therefore, we performed a micro-CT scan and three-dimensional reconstruction of the mouse femur at 6 weeks (Figure 8A, B). The results showed that the femur was basically intact, and there were no gaps after WKYMVm treatment; however, the PBS group showed obvious defects, followed by the WKYMVm+clodronate liposome treatment group. Quantitative analysis revealed that WKYMVm obviously increased the bone volume/total volume (BV/TV) compared to that in the other two groups (Figure 8C).

**Figure 8.**
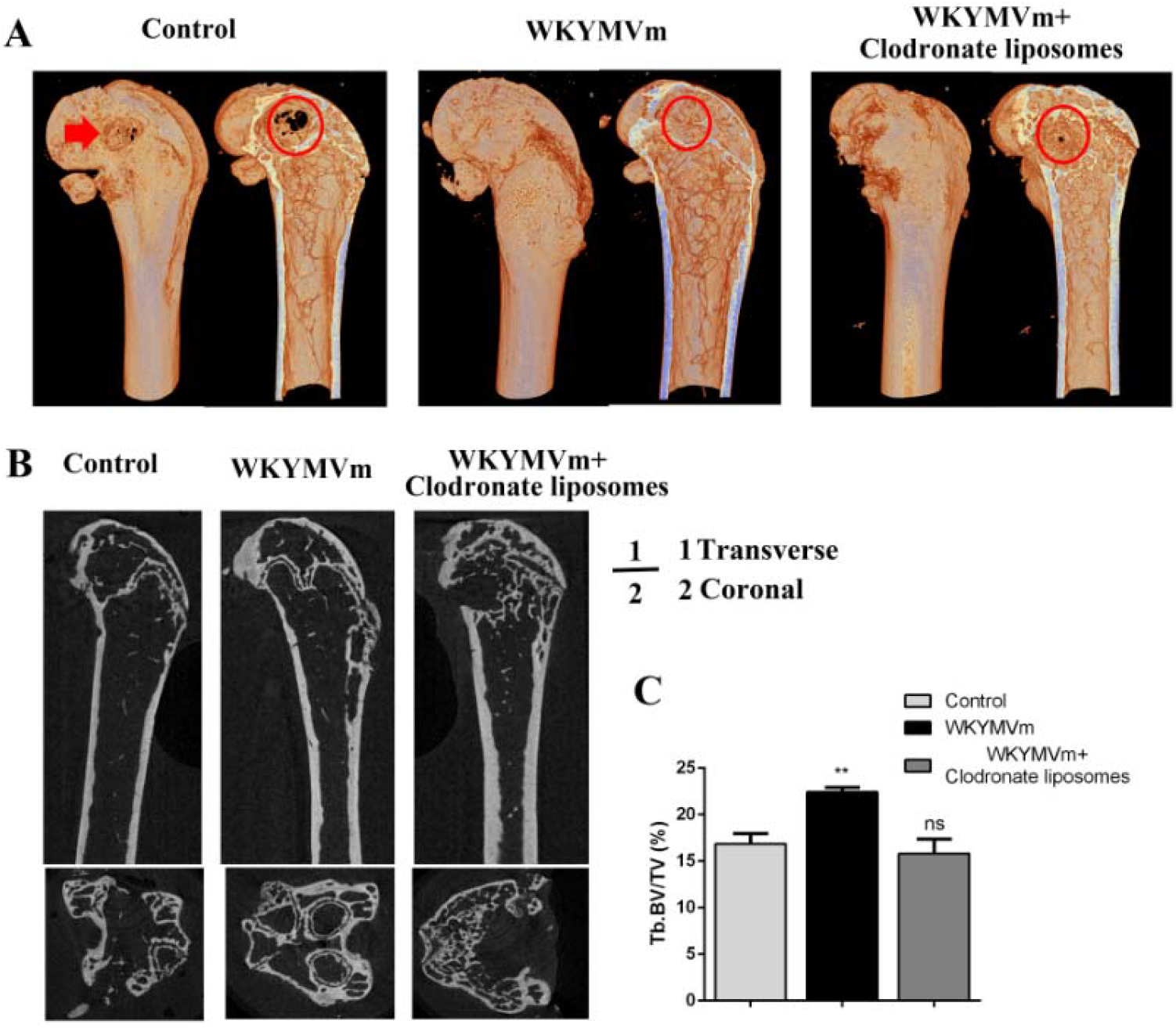
The combination treatment with WKYMVm promoted bone regeneration in vivo. A, Representative 3-dimensional reconstruction rendering at Week 6 from each experimental group to analyze bone formation at the defect site. B, Representative micro-CT images of distal femurs from mice receiving different treatments. C, Quantification of the percentage of bone volume/total volume (BV/TV). Data represent the means ± SD (n=3). *p < 0.05; **p < 0.01; and ***p < 0.001; ns, not significant relative to control.

## 4. Discussion

Vascularization is very important for the repair of bone defects. It has been reported that macrophages can differentiate into the proangiogenic M2 type in response to the cytokine environment. Therefore, the manipulation of macrophages to differentiate into an M2 phenotype holds great promise for bone repair. The current study found for the first time that the WKYMVm peptide can polarize macrophages into the M2 phenotype by secreting PDGF-BB through the JAK1/STAT6 pathway, which resulted in increased angiogenesis. The in vivo results confirmed that the enhanced M2/M0 ratio was mediated by WKYMVm administration, which promoted angiogenesis and repair of bone defects.

In our study, macrophage polarization to M2 but not to M1 was induced when RAW264.7 cells and/or BMMs were treated with WKYMVm (Figure 2). WKYMVm reportedly exhibits immunoregulatory ability, and its anti-inflammatory effect is widely recognized (30). Herein, it was speculated that WKYMVm may also be an effective agonist for macrophage immune regulation. This study first confirmed that WKYMVm-treated macrophages show low levels of iNOS, IL-1β, and TNF-α expression but high levels of Arg-1, TGF-β, and IL-10 expression in vitro (Figure 3), which is a characteristic of M2 polarization (12, 13), and M2 macrophages with this characteristic have been reported to have typical anti-inflammatory repair effects (14). Several other studies support our findings that WKYMVm activates different immune cells and modulates inflammatory activity (29, 30, 36).

The conditioned medium of WKYMVm-stimulated macrophages could better promote lumen formation in vitro (Figure 4C-G). In combination with our previous research, our findings indicate that WKYMVm induced M2 macrophages accompanied by increased PDGF-BB expression (Figure 4A). Similarly, ELISA results showed that PDGF-BB and VEGF secretions consistently increased with time. Interestingly, the increase in VEGF secretion stopped upon stimulation with 10 μM WKYMVm peptide for 48 h, at which time PDGF-BB secretion reached its maximum level (Figure 4B). The reason for this phenomenon is that the addition of WKYMVm increased VEGF secretion, with an increasing number of macrophages differentiated into the M2 phenotype, and PDGF-BB secreted by M2 cells plays a dominant role in the system. In addition, the secretion of PDGF-BB in the system was consistently significantly higher than that of VEGF. Therefore, it can be concluded that WKYMVm-induced M2 macrophage polarization plays a major role in pro-angiogenesis, resulting in the increased secretion of PDGF-BB to further promote angiogenesis.

Our study demonstrated that WKYMVm can regulate M2 macrophage polarization through JAK1/STAT6. Previous studies have widely confirmed that WKYMVm first highly efficiently binds to FPR2 on the membrane of monocytes and then transmits signals into the cells, thereby activating downstream proteins to exert their biological activities (37). In other words, FPR2 is an intracellular switch for WKYMVm signaling (36), and our results also confirmed that FPR2 is first activated in macrophages exposed to WKYMVm (Figure 5A). JAK1/STAT6 is a classic way to achieve polarization of M2 macrophages. Recently, it has been reported that WKYMVm can activate this pathway in human lung cancer cells, CaLu-6, to induce cell proliferation (28). However, it remains unknown whether this pathway is activated in macrophages. In this study, we wanted to determine whether WKYMVm works through the JAK1/STAT6 signaling pathway to regulate the polarization of M2 macrophages. The results showed that the total JAK1 and STAT6 expression in WKYMVm-stimulated macrophages did not change, but the JAK1 and STAT6 phosphorylation levels were significantly increased (Figure 5B). This is consistent with previous research showing that in immunomodulation and immune-mediated diseases, activation of STATs is closely related to signal activation of JAKs (38). A review article by O’Shea JJ showed that JAK-STAT signaling pathways are critically involved in various immune responses in immune cells through the action of hormones, interferons (IFNs), growth factors, and interleukins (39). In addition, PPAR-γ is an important transcription factor for macrophage polarization, and its activation is often accompanied by the activation of STAT6, which then co-induce M2 gene expression. Our data showed that the expression of PPARγ was enhanced significantly after WKYMVm stimulation (Figure 5D). Therefore, it can be concluded that WKYMVm activates the JAK1/STAT6 pathway. Furthermore, the addition of the JAK1 inhibitor ruxolitinib inhibited the expression of CD206 and Arg-1 in WKYMVm-induced macrophages (Figure 6A-C). Arg-1 is highly expressed upon STAT6 phosphorylation. Therefore, blocking JAK1 will prevent WKYMVm-induced macrophages from transforming into the M2 type, and at the same time, their angiogenic function will also be inhibited (Figure 6D-G). These results indicate that activation of JAK1/STAT6 by WKYMVm is necessary to induce M2 polarization and M2 functions. To our knowledge, this is the first report on the role of JAK1/STAT6 in the interaction between macrophages and WKYMVm. However, this study still has some shortcomings; we did not determine in detail the conditions to activate JAK after WKYMVm binds to the FPR2 membrane receptor, which needs further clarification in future research.

In a mouse femoral borehole defect model, WKYMVm induces M2 macrophages and the formation of new blood vessels, and this process is almost completely inhibited by macrophage-targeting clodronate liposomes. Furthermore, CD31 is an angiogenic marker, and a large number of studies have shown that adequate formation of new vessels could increase bone mass and promote bone regeneration (40). Our results agree with these reports; in the WKYMVm-induced group, the number of CD31+ vessels was the highest (Figure 7), and bone repair was the best (Figure 8). The remarkable plasticity of macrophages makes them interesting targets for immune regulation and tissue regeneration. The early inflammatory response in bone healing chemotactically attracts and recruits various repair cells; among them, macrophages are the key regulators of inflammation-related immunity (41). Recently, it has been found that the physiological inflammatory response is the main regulator of angiogenesis. Macrophages, especially M2 macrophages, can not only regulate inflammatory reactions but also promote angiogenesis to repair bone (19, 42). Our research supports the view that regulating the appropriate macrophage phenotype can promote functional recovery in bone defect repair and remodeling. The ability of M2 macrophages to promote angiogenesis in vivo in nontumor environments has also received increasing attention from researchers. In the field of orthopedics, cardiovascular disease, and other therapeutic fields, manipulating this type of macrophage with great plasticity has become a new treatment strategy (42–44). Here, we found that WKYMVm has immunomodulatory advantages, which provides additional evidence supporting the regulation of macrophages to promote angiogenesis at bone defect sites.

In summary, this study showed that WKYMVm regulates the polarization of M2 macrophages through the JAK1/STAT6 signaling pathway, thus promoting angiogenesis and the repair of bone defects. This study may provide alternative options for the regulation of macrophage phenotype and function, and WKYMVm appears to have the potential to immunomodulate bone defects.

## Acknowledgments

This work was supported by grants from the National Natural Science Foundation of China (Grant No. 81672177), the Southwest Hospital Clinical Innovation Foundation (Grant No. SWH2016JCZD-05, SWH2016JSTSZD-05, SWH2014 LC-20).

## Author contributions

X. Han and J. Hu completed the in vitro studies; X. Han and W. Zhao performed the in vivo studies; X. Han and J. Dai analyzed the data; X. Han wrote the paper; X. Han and Q. He designed the study; and X. Han and Q. He prepared the final manuscript.

## Data Accessibility

The data used to support the findings of this study are available from the corresponding author upon reasonable request.

